# Transcriptional reprogramming of natural killer cells by vaccinia virus shows both distinct and conserved features with mCMV

**DOI:** 10.1101/2022.11.10.516015

**Authors:** Delphine M Depierreux, Geoffrey L Smith, Brian J Ferguson

## Abstract

Natural killer (NK) cells have an established role in controlling poxvirus infection and there is a growing interest to exploit their capabilities in the context of poxvirus-based oncolytic therapy and vaccination. How NK cells recognise poxvirus-infected cells to become activated remains unclear. To address this knowledge gap, we studied the NK cell response to vaccinia virus (VACV) *in vivo*, using a systemic infection murine model. We found broad alterations in NK cells transcriptional activity in VACV-infected mice, consistent with both direct target cell recognition and cytokine exposure. There were also alterations in the expression levels of specific NK surface receptors (NKRs), including the Ly49 family and SLAM receptors, as well as upregulation of memory-associated NK markers. Despite the latter observation, adoptive transfer of NK memory populations did not confer protection from re-infection. Comparison with the NK cell response to murine cytomegalovirus (MCMV) infection highlighted common features, but also distinct NK transcriptional programmes initiated by VACV. Finally, there was a clear overlap between the NK transcriptional response in humans vaccinated with an attenuated VACV, modified vaccinia Ankara (MVA), demonstrating conservation between the NK response in these different host species. Overall, this study provides new data about NK cell activation, function, and homeostasis during VACV infection, and may have implication for the design of VACV-based therapeutics.

## Introduction

Natural killer (NK) are effector lymphocytes that form part of the innate immune system and function in the clearance of stressed cells such as virus-infected or tumour cells (Vivier et al., 2008). During virus infection, NK cells deploy cytotoxic and cytolytic functions to clear infected cells and secrete immune-regulatory effectors that shape the subsequent immune response. NK cells can also mature to develop memory-like capabilities. A large body of evidence links NK cell function to the control and clearance of viruses including human immunodeficiency virus (HIV), hepatitis C virus (HCV), SARS-CoV-2, influenza virus, poxviruses and herpesviruses (Biron et al., 1999; Jost and Altfeld, 2013; Maucourant et al., 2020). This effect is most starkly indicated by individuals with primary immune deficiencies, such as X-linked lymphoproliferative disease, which result in the loss of NK cells or NK functions, and cause high susceptibility to repeated virus infections (Orange, 2013).

NK cells control viral spread by killing virus-infected cells by direct cytolysis, secreting immunoregulatory molecules, such as IFNγ, CCL4 and TNFα, which attract and regulate the activation of other immune cells. In some cases, NK cells also develop a memory-like response. NK cells rely on the expression of surface receptors, collectively called NKRs, to sense inflammation and recognise abnormal cells. These receptors provide a balance of activating and inhibitory signalling in the presence of a healthy cell. This balance is perturbed when there are abnormal surface protein expression patterns on neighbouring cells, leading to NK activation. By sensing a variety of stimuli, an appropriate timing and degree of activation is achieved (Lanier, 2008; Vivier et al., 2008) that allows carefully regulated killing of infected cells and avoids unnecessary tissue damage. Activating NKRs detect ligands such as viral molecules like the m175 cell surface protein from murine cytomegalovirus (mCMV) and the influenza virus haemagglutinin, and cellular stress-induced ligands such as human MICA/B and ULBP and mouse RAE1 (Arase et al., 2002; Lee and Biron, 2010). Inhibitory receptors provide opposing signals and recognise constitutively expressed self-molecules such as MHC class I. Loss of MHC class I, for example on virus-infected cells, results in a reduction in inhibitory signalling to NK cells and contributes to their activation by recognition of missing-self (Ravetch, 2000). NK cells can also be activated by antibody-coated target cells via their expression of the low-affinity Fc receptor, CD16, providing a signal for ADCC. The expression of a range of receptors for inflammatory cytokines, such as IL-12, IL-15, IL-2 and IL-18, and type I interferons (IFNs), allows NK cells to sense and respond to the inflammatory environment produced by infection or tissue damage. NK cells then respond by maturation, proliferation and initiation of effector functions (Vivier et al., 2008; Costanzo et al., 2018). Using these multiple sensing mechanisms, NK cells contribute significantly to a rapid, early response to infection (Biron et al., 1999; Vivier et al., 2008; Jost and Altfeld, 2013). Subsequently, a subset of NK cells can take on memory-like properties, upregulating markers such as Thy1 and CXCR6 and contributing to recall response to infection and hapten challenge (Sun et al., 2009a; Gillard et al., 2011).

Vaccinia virus (VACV) was used as the vaccine to eradicate smallpox. Attenuated forms, such as modified vaccinia Ankara (MVA), are currently in use as vaccine against monkeypox (Earl et al., 2004) and in clinical trials for heterologous vaccine vectors and oncolytic therapeutics (Smith et al., 2013). NK cells have an established role in controlling multiple poxvirus infections (Jacoby et al., 1989; Parker et al., 2007) and the depletion of NK cells increases susceptibility to VACV (Bukowski et al., 1983). In addition, VACV infection induces proliferation and accumulation of NK cells at the site of infection (Natuk and Welsh, 1987; Dokun et al., 2001; Jacobs et al., 2006) and VACV infection increases cell susceptibility to NK lysis *ex vivo* (Brutkiewicz et al., 1992; Baraz et al., 1999; Chisholm and Reyburn, 2006). Multiple studies have reported that NK cells require the presence of lL-12 and IL-18 (Gherardi et al., 2003; Brandstadter et al., 2014), and type I IFN to control VACV infection (Martinez et al., 2008). Additionally, following VACV infection, a subset of Thy1-expressing memory-like NK cells was reported to contribute to protection against homologous challenge (Gillard et al., 2011). There is little data, however, about how NK cells become activated, which of their receptors are engaged in the recognition of VACV-infected cell and how NK cells are reprogrammed in response to VACV infection.

To address this knowledge gap, we analysed changes in the transcriptome of circulating NK cells following intranasal infection with VACV strain Western Reserve (WR), a model in which the virus can spread from the respiratory system to other organs (Williamson et al., 1990; Alcamí and Smith, 1992). These changes were then compared to similar NK transcriptomic datasets from humans infected with MVA and mice infected with mCMV. We also compared the NK transcriptional profile with the abundance of cell surface molecules determined by flow cytometry. We show that VACV infection induces activation, expansion and maturation of NK cells with concomitant changes in the bulk transcriptional program and expression of memory markers. The transcriptional response is consistent with NK cells becoming activated by a combination of direct recognition of infected cells and by exposure to cytokines and IFN. Comparison with NK transcriptional responses to MVA vaccination in humans and in response to different viruses, including mCMV, indicates conserved features of NK transcription reprogramming in the context of vaccination or infection.

## Materials and Methods

### Mouse husbandry

All animal experiments were conducted according to the Animals (Scientific Procedures) Act 1986 under the license PPL 70/8524. Mice were housed in specific pathogen-free conditions in the Cambridge University Biomedical Services facility. Female C57BL6 mice, 6-8 weeks old from Charles River were used throughout the study.

### *In vivo* infection models

For the intranasal (i.n.) model, mice were anaesthetized with isoflurane and inoculated with 5 × 10^3^ plaque-forming units (p.f.u.) of wild type (WT) VACV strain WR for primary infection, or with 10^5^ p.f.u for post-vaccination challenge, or were injected with vehicle control. For the intradermal (i.d.) infection model, mice were anaesthetized with isoflurane and infected i.d. in the ear pinnae with 10^3^ p.f.u. of WT VACV strain WR. For all infections, VACV strain WR was purified from cytoplasmic extracts by sedimentation through a sucrose cushion. Inocula were prepared in phosphate-buffered saline (PBS) supplemented with 0.01% bovine serum albumin (BSA, Sigma Aldrich) and the infectious titres administered were confirmed by plaque assay on BSC-1 cells.

### Adoptive transfer

Four weeks post i.d. infection, spleens and blood were collected under sterile conditions from mice. Blood was collected by heart puncture into microvette CB 300 μl tubes with clot activator (Sarstedt) and left at RT for 2 h to allow clot formation. Serum was isolated after centrifugation at 10,000 *g* for 5 min at RT. Spleens were processed as described below to isolate NK or CD8+ T cells. Recipient mice were injected intravenously with NK cells, CD8+ T cells or splenocyte suspensions in the tail veil and challenged i.n. with 10^5^ p.f.u. of VACV WR 24 h later. Mice were weighed daily and monitored for signs of illness for two weeks post challenge.

### Flow cytometry

Mouse spleens were disrupted mechanically through a 70-µm cell strainer (Falcon, BD) to obtain a single-cell suspension. Erythrocytes were lysed with 2 ml of lysis buffer (BD Pharmlyse) according to manufacturer’s instructions. PBS was added to quench the lysis reaction and samples were centrifuged at 500 *g* for 5 min. Cells were then counted using a Nucleocounter NC-250 (Chemometec). Subsequently, single-cell suspensions were labelled for phenotypic analysis or sorting. Cells were labelled with Zombie viability dye (Biolegend) for 15 min at 4 °C. Fc receptors were blocked with 10 μg/ml anti-mouse CD16/CD32. Abs for surface markers (Table 1) were added for 30 min at 4 °C in the dark. Cells were washed with PBS twice and collected by centrifugation at 300 *g* for 5 min at 4 °C. The relevant fluorescence-minus-one (FMO) and isotype mAb labelling conditions were included as controls. All samples were fixed with 4 % paraformaldehyde (PFA) before acquisition by flow cytometry on an LSR Fortessa (BD).

**Table 1.**
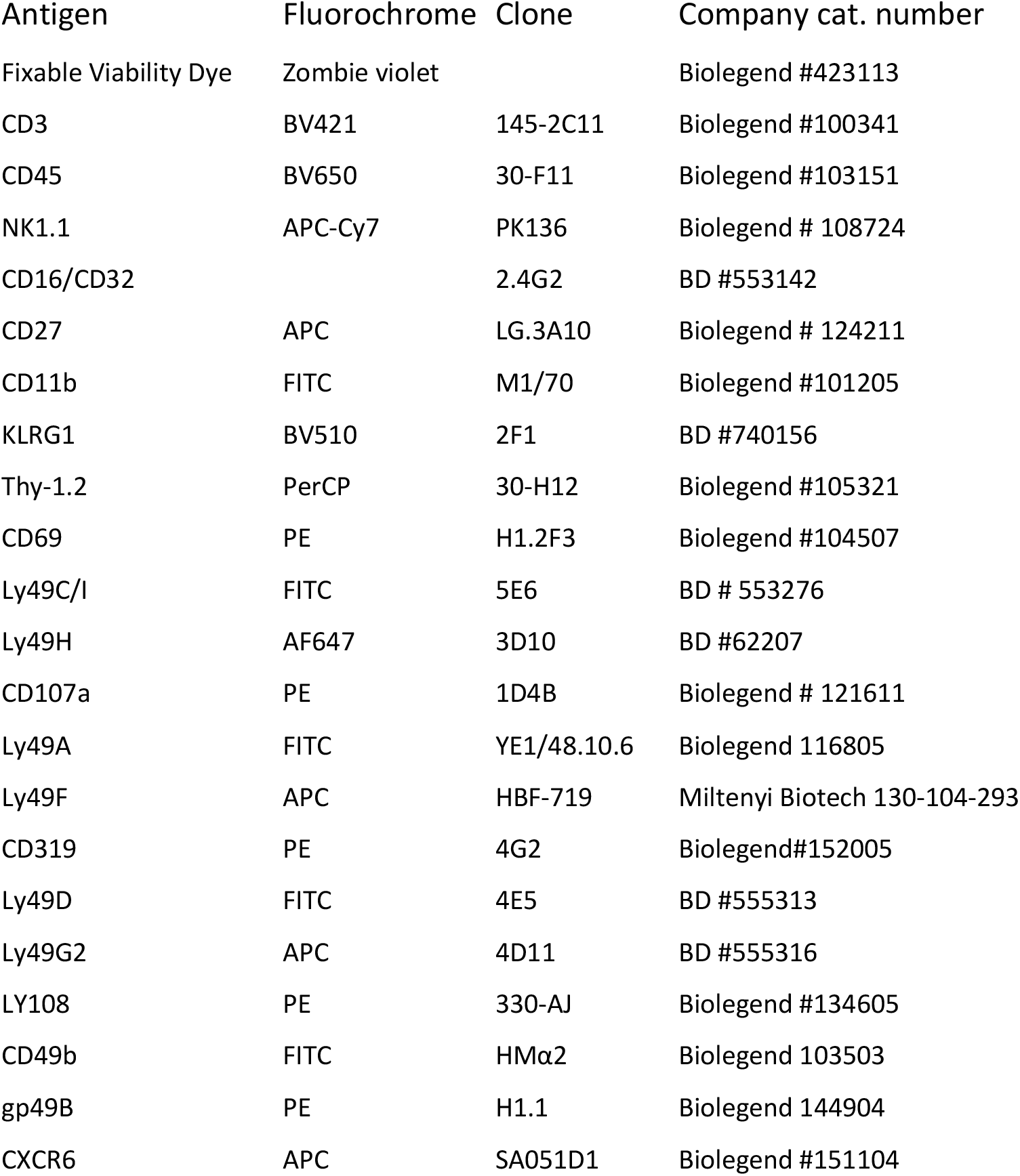
Antibodies used for Flow cytometry

**Table 2.** Number of genes differentially expressed for the indicated pairwise comparisons during RNA-seq study of murine NK cells. Groups of B6 mice (n=4) were i.n. infected with WT VACV or vehicle control for 1.5 or 6.5 d.p.i‥ Differential expression of transcripts was analysed following 4 pairwise comparisons, as indicated in the first column. The number of transcripts differentially expressed in a statistically significant manner is indicated in the second column (FDR<0.01) and in the third column (FDR<0.0.5). The total number of genes detected is indicated in the last column.

| Comparison | FDR < 0.01 genes | FDR < 0.05 genes | Total genes |
| --- | --- | --- | --- |
| Mock_D6.5__VS__WT_D6.5 | 2314 | 3280 | 9647 |
| Mock_D1.5__VS__WT_D1.5 | 16 | 70 | 9646 |
| Mock_D1.5__VS__Mock_D6.5 | 55 | 140 | 9664 |
| WT_D1.5__VS__WT_D6.5 | 2977 | 4045 | 9645 |

### Lymphocyte isolation

For NK cell isolation, splenic single cell suspensions were resuspended in 600 µl PBS, and Fc receptors were blocked with 10 μg/ml anti-mouse CD16/CD32. Cells were incubated with viability dye (Zombie Violet, BioLegend) for 15 min in the dark. Cells were washed twice in PBS and a cocktail of 10 µg anti-CD3-APC, 10 µg anti-NK1.1-PE and 50 µg anti-CD45-FITC (Supplementary Table 1) was added to each sample. After 20 min of incubation in the dark, cells were washed twice in PBS, collected by centrifugation, and resuspended in 1.5 ml PBS. NK cells were sorted by flow cytometry. A sample of sorted NK cells was rerun on the flow cytometer to assess purity. CD8+ T cells were isolated using a CD8a+ T Cell Isolation Kit (Milteny, 130-104-075). The purity and the presence of VACV-specific CD8+ T cells were assessed by staining isolated CD8+ T cells as described above with zombie violet viability dye, CD8-APCR700, CD3-BV605, and vaccinia virus B8 Dextramer-PE (Immunodex) following the manufacturer’s instructions.

### Bulk RNAseq

Half a million viable NK cells (CD45^+^CD3^−^NK1.1^+^ cells) (Supplementary Figure 1A) from spleens of mock and VACV-infected mice were sorted by FACS in sterile PBS. NK cell purity was >96 % (Supplementary Figure 1B). Directly after cell sorting, the samples were centrifuged at 7500 *g* for 7 min. The pellets were resuspended in 350 µl of RLT buffer (Qiagen) supplemented with β-mercaptoethanol (10 µl/ml). Samples were snap-frozen and stored at −80 °C until RNA extraction.

Total RNA was extracted from purified NK cells using the RNeasy Mini Kit (Qiagen) with an additional DNAse treatment (Qiagen RNase-Free DNase Set). RNA concentration, purity and integrity were checked using a SpectroStar FLUOstar OMEGA (BMG LabTech) and an Agilent 2100 Expert bioanalyser (Agilent Technologies). The SMART-Seq v4 Ultra Low Input RNA Kit for Sequencing (Takara Bio) was used to generate cDNAs and quality was assessed using an Agilent 2100 bioanalyser. The Nextera XT DNA Library Prep Kit (Illumina) was used to generate libraries from 125 pg of input cDNA. Library quality was checked with a tapestation (HS D1000) and a Qbit with dsDNA high sensitivity. Libraries were normalised to the lowest concentration and pooled together to be run on the NextSeq500 (Illumina) for single-end sequencing on 75 cycles.

For differential expression analysis, the threshold was set to include genes with at least 5 counts per million reads mapped and detected in at least half the samples to allow for on/off expression pattern recognition. Normalisation factors were calculated by considering the sequencing depth and RNA composition. A correction was applied for G+C content and gene length bias by using CQN Bioconductor package v.1.24.0 that allows removal of a systematic G+C content and gene length bias by a smoothing function. Pairwise comparisons between groups were performed using the counted reads and the R package edgeR version 3.16.5. The resulting p-value was then corrected for multiple hypothesis testing using the Benjamini-Hochberg method.

### Biological process gene ontologies enrichment

To assess the biological processes affected during VACV infection in NK cells, significant (FDR <0.05) differentially-expressed genes (DEG) between mock and VACV-infected samples were analysed in R using BioConductor. DEG lists were analysed for biological process GO (Gene Ontology) enrichment against a background made of all the genes detected in the mock and WT-infected samples. The gseGO function was used for gene set enrichment analysis (p value cut-off 0.05, minimum gene set size 10, maximum gene set size 500). The simplify function was used to aggregate redundant GO. Enrichment scores (p-values) were corrected for multiple hypothesis testing using the Holm-Bonferroni method. The most significant GO were plotted using the dotplot function.

### Software

GraphPad Prism 4.0 and R3.0 were used for statistical analyses and graphic representation. FlowJow Tree Star was used for flow cytometry data analysis.

## Results

### NK cells undergo transcriptional reprogramming following VACV infection

To probe the NK cell response to VACV infection, C57BL/6 mice were infected i.n. with VACV strain WR. An expansion of NK cell numbers was observed at 6.5 d post infection (d.p.i.) (Figure 1A), the time at which the NK response to VACV is reported to peak (Jacobs et al., 2006; Abboud et al., 2016). The relative expression of CD27 and CD11b (Chiossone et al., 2010) defines maturation subsets in NK cells and the analysis of these markers at 6.5 d.p.i. by flow cytometry indicated preferential expansion of CD27+CD11b+ and CD27+CD11b-subsets both in terms of absolute numbers and as a percentage of total NK cells (Figure 1B, C). These data are consistent with the preferential engagement of NK subsets that are potently cytolytic and have the capacity to secrete high levels of cytokines.

**Figure 1:**
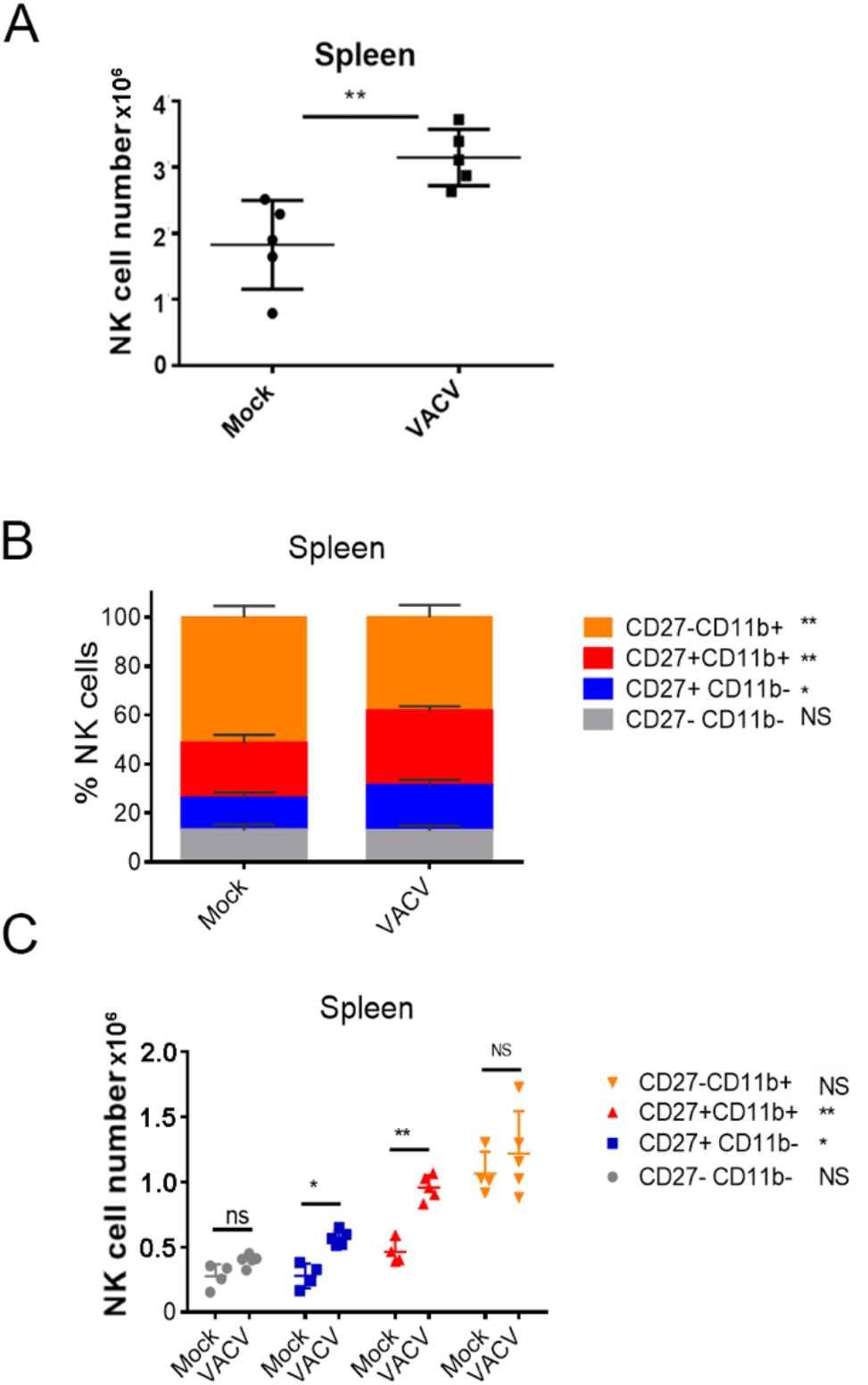
Expansion and maturation of splenic NK cells in response to VACV infection. C57BL/6 mice were mock-treated or infected i.n. with VACV and 6.5 d later (A) splenic NK cells were counted. (B, C) splenic NK subsets were quantified by flow cytometry for CD27 and CD11b expression. Data presented are representative of two independent experiments n=5 *p<0.05, **p<0.01.

To define the alterations in the transcriptional programme of NK cells during VACV infection we isolated populations of splenic NK (CD45+CD3-NK1.1+) cells at 1.5 and 6.5 d.p.i. and generated a transcriptional profile by bulk RNAseq. We chose 1.5 and 6.5 d.p.i. to understand alterations in NK phenotype at early and peak times post VACV infection. Differential gene expression analysis between VACV-infected and their respective mock samples, at 1.5 and 6.5 d.p.i. revealed, respectively, 70 and 3280 transcripts significantly altered (FDR<0.05). Pairwise comparisons of the two VACV-infected samples with each other revealed 4045 significantly altered transcripts. Comparison between the two mock samples, revealed 140 transcripts significantly altered (FDR<0.05) (Supplementary Table 2).

Hierarchical clustering showed that all 4 VACV-infected mice samples at 6.5 d.p.i. clustered together, away from the other 12 samples (Figure 2A). The 4 VACV-infected samples at 1.5 d.p.i. (WT-D1.5-m) clustered together with the eight mock samples, consistent with the number of genes that were differentially expressed (Supplementary Table 2). This suggests that differentially expressed genes (DEGs) in WT infected mice at 1.5 d.p.i. are indistinguishable from background noise, whilst those detected in infected mice at 6.5 d.p.i. are biologically different in response to VACV infection. Analysis of the DEGs between VACV-infected and mock-infected mice showed that the most significantly altered transcripts included multiple NK cell receptors (NKRs) (*Gp49A/B*, T cell immunoreceptor with Ig and ITIM domains (*Tigit*)), effector molecules (*Serpinb9b, Gzmk, Serpin3af*), IFN-stimulated genes (ISGs) (*Gbp6, Ifi44, Gbp10, Stat1*) and cell cycle regulators (*Cables1*) (Figure 2B). Furthermore, multiple biological processes were significantly enriched in VACV-infected samples at 6.5 d.p.i. compared to mock (FDR<0.05) (Figure 2C). These suggest that NK cells display an active defence phenotype and engage effector mechanisms in response to VACV infection.

**Figure 2.**
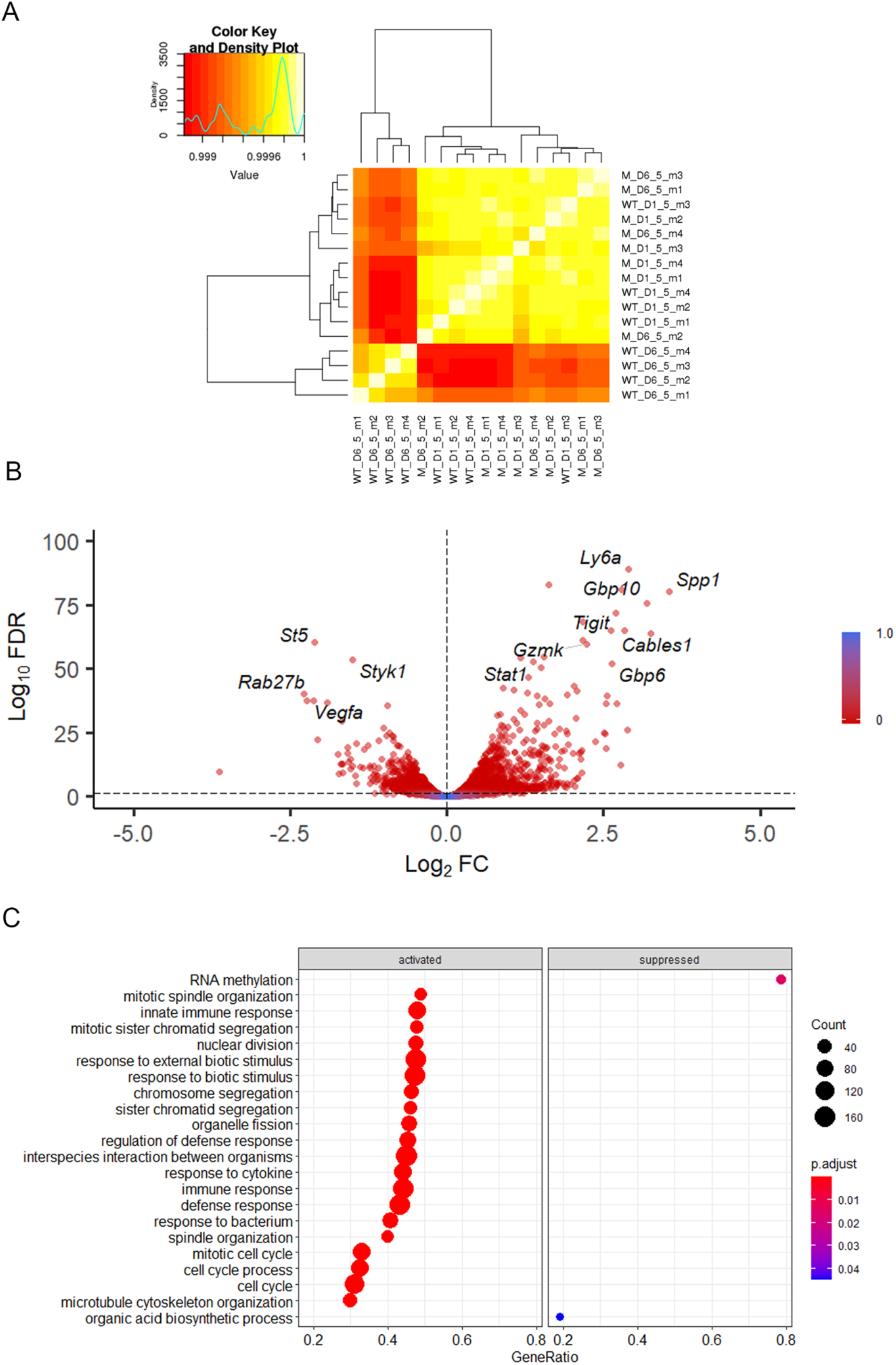
Transcriptomic analysis of splenic NK cells following VACV infection. C57BL/6 mice (*n*=4) were infected i.n. with WT VACV or were mock-infected with vehicle control for 1.5 or 6.5 d and the transcriptome of splenic NK cells was analysed by RNA-seq. (A) Heatmap representing correlations among the 16 RNA-seq samples. The intensity of the cell colour corresponds to the degree of correlation between the compared groups. “WT” represents infected mice, “M” the mock-infected mice. (B) Volcano plot of NK cell transcripts differentially expressed at 6.5 d.p.i. in VACV-infected mice compared to mock. Each DEG (FDR<0.05) (WT vs mock) is represented by a blue dot. (C) Biological process enrichment amongst murine NK cell transcripts upregulated during VACV infection. Upregulated (FDR<0.05) transcripts.

### VACV infection activates NK cell effector mechanisms

Combining further analysis of the NK cell transcriptome (Figure 3A, B) with targeted flow cytometric quantification of cell surface markers (Figure 3C), showed significant upregulation of transcripts and proteins associated with NK cell proliferation, activation and effector functions. This included *Il2r* and *Ki67*, two proliferation markers, as well as granzymes, IFNγ and TNF superfamily member genes, which are indicative of increases in effector functions (Figure 3A). NK cell surface activation markers (CD69, KLRG1, CD107a, GP49b) showed significant upregulation measured by flow cytometry (Figure 3C). The transcripts encoding these proteins were also upregulated, except from *Lamp1* (CD107a), indicating a strong correlation between mRNA and protein levels and suggesting a significant transcriptional contribution to NK cell activation following VACV infection (Figure 3B).

**Figure 3.**
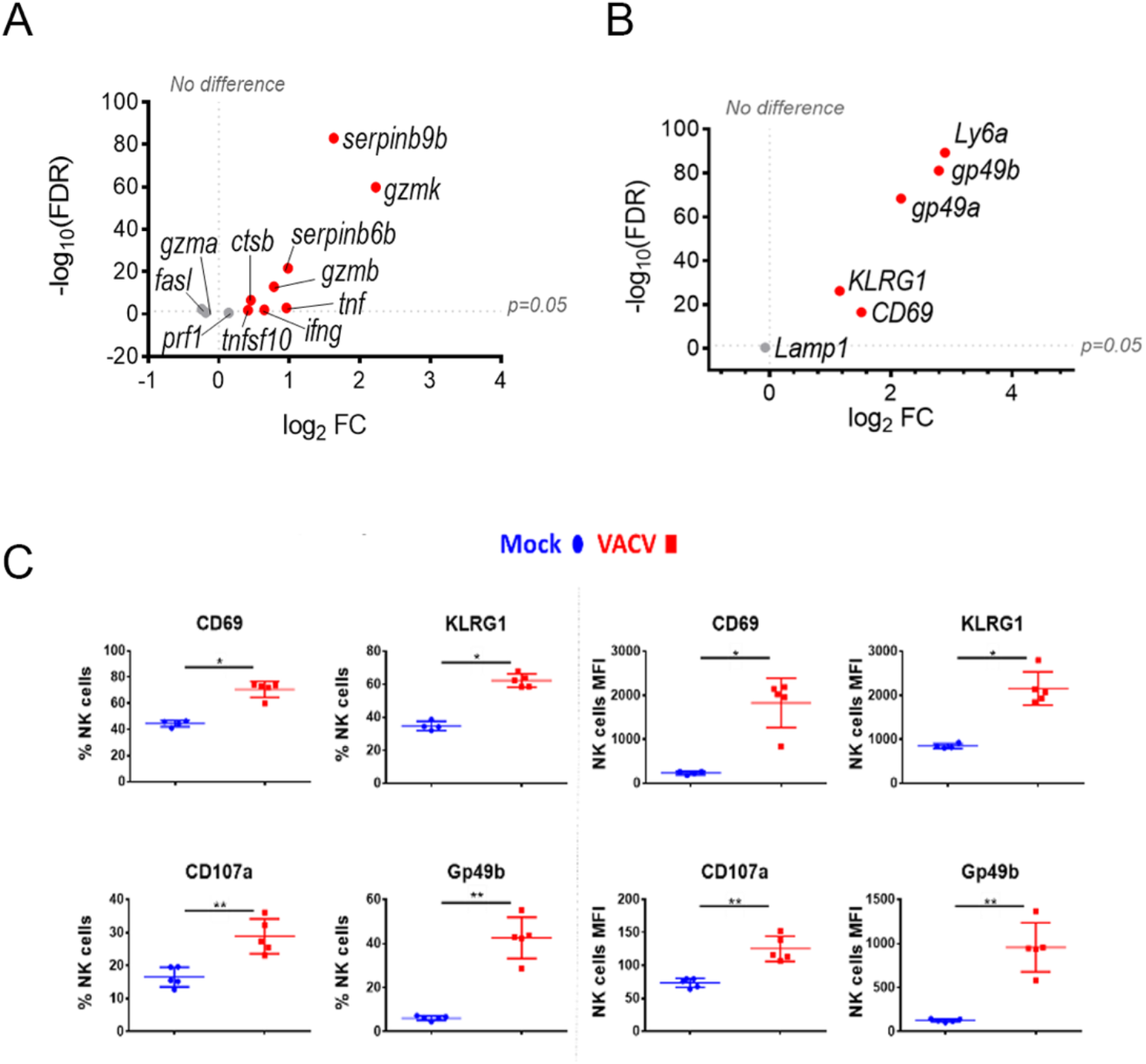
NK cells become activated and upregulate key effector mechanisms during VACV infection. C57BL/6 mice were infected i.n. with VACV or were mock-infected with vehicle control for 6.5 d and then NK cells were isolated for bulk RNA-seq (n=4) or flow cytometry (n=5). (A) The differential expression of effector proteins transcripts (VACV vs mock) was analysed by RNA-seq. Each dot represents a transcript. Red indicates statistical significance (FDR<0.05). (B) Quantification of NK cell surface activation markers in mock (blue) and VACV-infected mice (red) by flow cytometry. NK cells expressing the indicated marker by percentage and MFI are shown. Error bars represent ± SD. Statistical significance was calculated using a Mann-Whitney test (*p<0.05, ** p<0.01, ***p<0.001). (C) Differential expression of activation marker transcripts (VACV vs mock). Each dot represents a transcript and red indicates statistical significance (FDR<0.05).

To discover which stimuli contribute to NK cell activation during VACV infection *in vivo*, our RNA-seq data were compared with a published study describing the unique transcriptomic fingerprint of NK cells activated by ADCC, target cell recognition or cytokine (Costanzo et al., 2018). Gene sets corresponding to the unique NK cell transcriptomic signature associated with these 3 independent stimuli compared with our RNA-seq dataset. Among the upregulated transcripts, 48 (direct recognition), 42 (cytokine stimulation), and six (ADCC) transcripts were detected in our dataset. Side-by-side comparison of the expression pattern of such transcripts showed that our RNA-seq data matches primarily with the transcriptomic signature corresponding to direct recognition. Transcripts for this category presented the highest upregulation in our dataset (Figure 4A), whereas genes involved in response to cytokines were less modulated (Figure 4B), and genes involved in the ADCC response showed slight or no upregulation. In addition to this comparative analysis, a set of ISGs (*Ifi44, Ifi211, Irf2, Irf7, Ifit1, Ifit3, Gbp10* and *-6, Oas1, Cxcl10* and *ligp1)* was upregulated in our dataset (Figure 5) consistent with NK cells responding to IFN exposure. Overall, this analysis indicates that NK cells recognise abnormal protein expression pattern at the surface of VACV-infected cells *in vivo*, but also that cytokines and IFNs participate in this activation.

**Figure 4.**
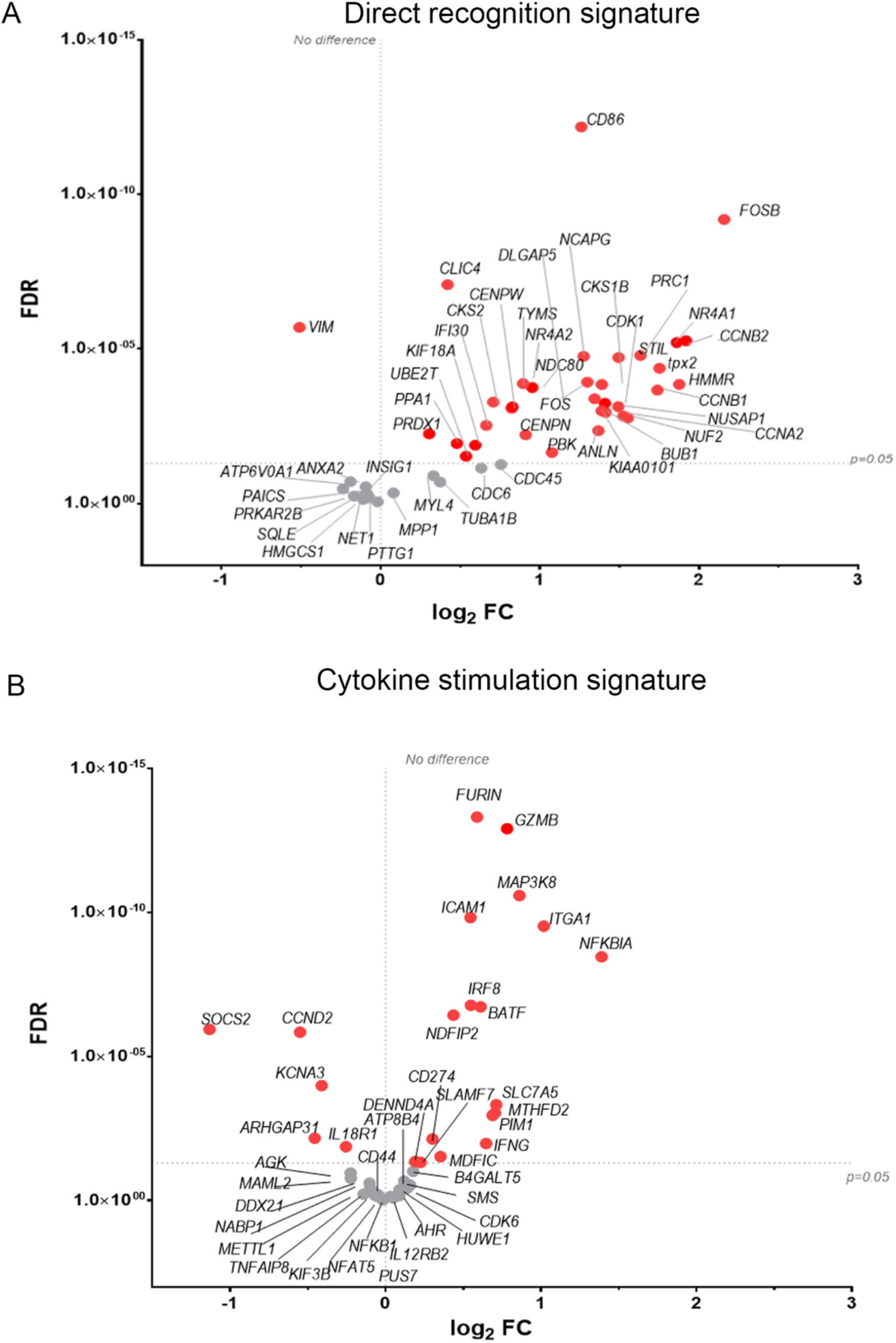
NK cells exhibit transcriptional signatures consistent with direct cell recognition and cytokine stimulation. The differential expression profile (VACV vs mock) of transcripts detected in RNA-seq that are indicative of (A) direct recognition of infected cells, or (B) cytokine stimulation are shown. Each transcript is represented by a dot, its FC (WT vs mock) is shown on the Y-axis, and statistical significance of the FC (FDR) is shown on the X-axis. Comparator dataset reference GSE110446. Red indicates statistical significance (FDR<0.05).

**Figure 5.**
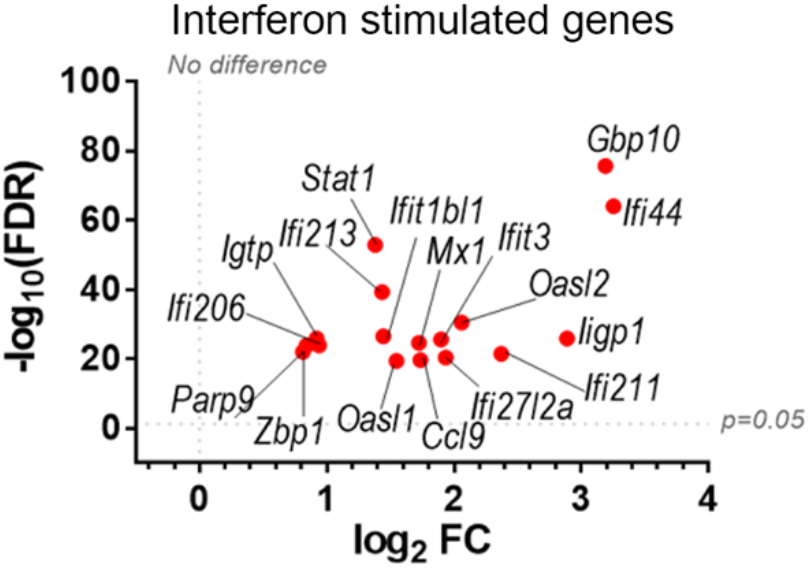
Upregulation of ISGs transcripts in NK cells from VACV-infected mice compared to mock. C57BL/6 mice (*n*=4) were infected with VACV WT or were mock-infected. At 6.5 d.p.i., NK cell transcriptional changes were analysed by RNA-seq, and differential expression (VACV vs mock) was calculated for each gene. Seventeen ISGs were found out of the 100 most significantly altered transcripts (VACV vs mock (FDR<0.05)) and are depicted here. Each dot represents the differential expression of an ISG transcript (WT vs mock). Red indicates statistical significance (FDR<0.05).

### VACV infection modulates NK cell receptor surface expression

Direct recognition of infected cells by NK cells is mediated by NKRs. Here, we analysed how NKR expression is modulated during VACV infection. A manually curated list of NKRs known to influence the activation status of NK cells was generated (Lanier, 1998, 2008; Ravetch, 2000; Kelley et al., 2005; Vivier et al., 2008) and their expression profile was analysed in our RNA-seq dataset (Figure 3). Several NKR transcripts were significantly upregulated (*tigit, gbp49a/b, klrg1, cd69, crtam, lag3, thy1, klrb1b, cd160 and lair1*) or downregulated (*klra9, klrc2*), whilst other were less affected. We combined these transcriptional data with flow cytometric analysis to reveal how specific families of NKRs are involved during VACV infection.

Receptors from the Ly49 family interact with MHC-I, a major NK ligand, and some Ly49 receptors engage directly with viral ligands (Arase et al., 2002; Smith et al., 2002; Kielczewska et al., 2009; Pyzik et al., 2011) and LY49H define mCMV-specific memory NK cells subsets (O’Leary et al., 2006; Sun et al., 2009b; Paust et al., 2010; Wight et al., 2018). Therefore, we analysed the expression of such receptors at the transcript (Figure 6A) and protein (Figure 6B) level in NK cells during VACV infection. Ly49H, Ly49D and Ly49G2 expression was not altered during VACV infection in mice (Figure 6) whilst Ly49C/I were downregulated, both at the protein and the transcriptomic level, and with a greater fold change (FC) for Ly49I (Figure 6). Further, our flow cytometry data showed significant upregulation for Ly49A both on splenic NK cells by flow cytometry and by analysis of Ly49A transcripts (Figure 6). Finally, our data showed that Ly49F transcripts and surface protein expression were upregulated in splenic NK cells (Figure 6).

**Figure 6.**
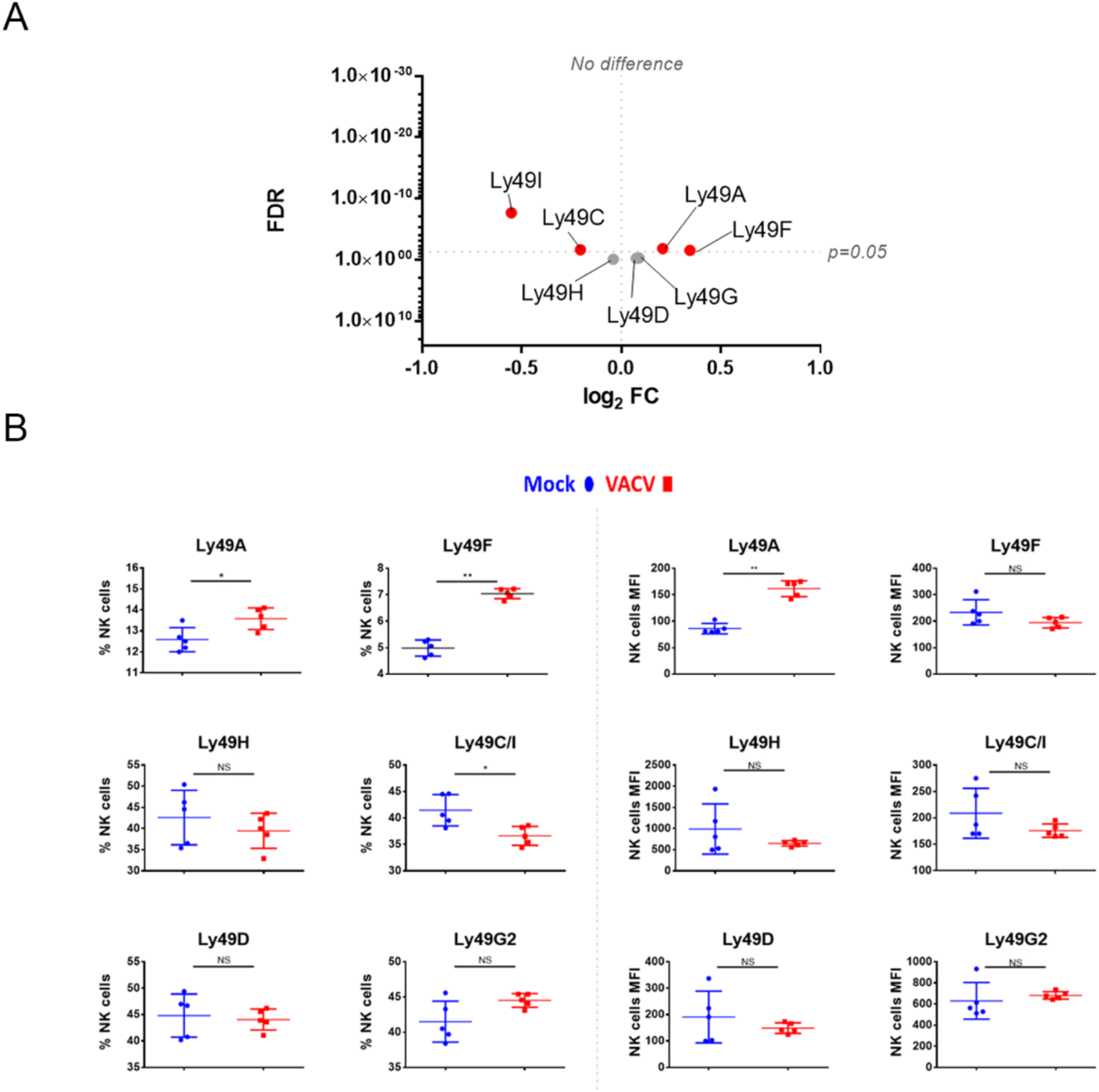
Differential expression of Ly49 family receptors at the transcriptional and protein level during VACV infection. (A) C57BL/6 mice (*n*=4) were infected with VACV WT or were mock-infected. At 6.5 d.p.i., NK cell transcriptional changes were studied by RNA-seq (*n*=4), and differential expression (VACV vs mock) was calculated. Each dot represents the differential expression profile of a Ly49 transcript (WT vs mock). Red indicates statistical significance (FDR<0.05). For clarity, the protein name is indicated rather than gene name. (B) Protein expression assessed by FACS for Ly49 receptors in splenic NK cells from mock (blue) and VACV-infected mice (red). The percentage of NK cells expressing the receptor and the MFI for the indicated receptor is shown. Error bars represent ± SD, statistical significance was assessed with a Mann-Whitney test (*p<0.05, ** p<0.01, ***p<0.001).

We next analysed SLAM receptor expression. CRACC (slamf7) and Ly108 (slamf6) were both upregulated in splenic NK cells at both at the transcript and protein level (Figure 7A, B). Transcripts for the activating adaptors EAT2A/B, SAP, and ERT were upregulated whilst transcripts of inhibitory mediators were downregulated or unchanged, suggesting an activating function for SLAM receptors. Additionally, the transcriptomic data showed that the expression of other SLAM receptors transcripts for 2B4 (slamf4), CD48 (slamf2) and CD229 (slamf3) was mildly downregulated or unaffected (Figure 7A). Taken together, these data indicate that CD319 and Ly108 are the two members of the SLAM family whose expression is upregulated in NK cells in the context of VACV infection. These receptors are likely to mediate activating functions via their co-expression with activating adaptors.

**Figure 7.**
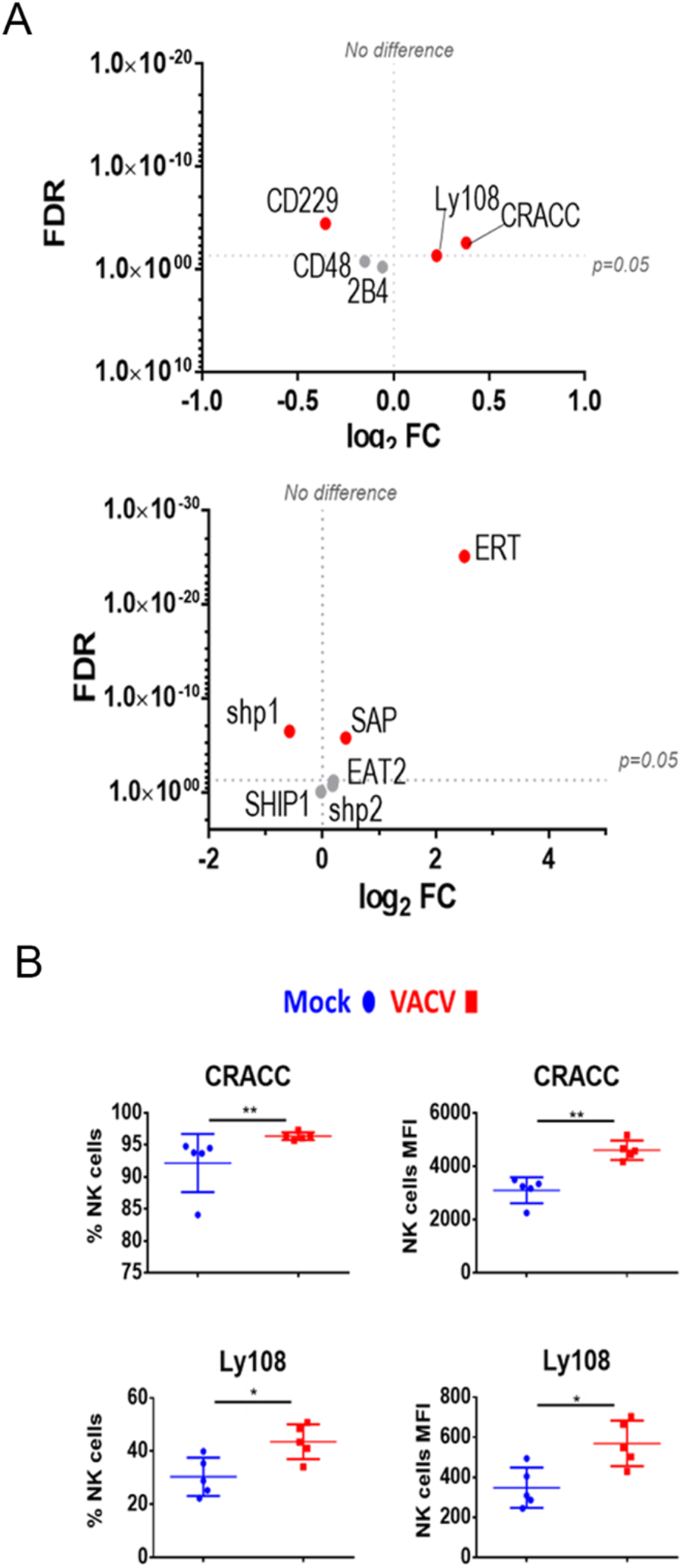
Differential expression of SLAM family receptors at the transcriptomic and proteomic levels in NK cells during VACV infection. (A) C57BL/6 mice (n=4) were infected with VACV WT or were mock-infected. At 6.5 d.p.i., NK cell transcriptional changes were studied by RNA-seq (n=4), and differential expression (VACV vs mock) was calculated. Each dot represents the differential expression profile of a SLAM family receptor or adaptor protein transcript (WT vs mock). Red indicates statistical significance (FDR<0.05). For clarity, the protein name is indicated rather than gene name. (B) Protein expression assessed by FACS for SLAM receptors in splenic NK cells from mock (blue) and VACV-infected mice (red). The percentage of NK cells expressing the receptor (prevalence) and MFI (expression level) for the indicated receptor is shown. Error bars represent ± SD, statistical significance was assessed with a Mann-Whitney test (*p<0.05, ** p<0.01, ***p<0.001).

### VACV infection leads to upregulation of transcripts associated with memory development

Markers of NK memory such as Thy1 and CXCR6 (Paust et al., 2010; Gillard et al., 2011), were also analysed. Both markers were significantly upregulated at the transcriptomic level and Thy1 protein level was upregulated (Figure 8A, B). Other NK cell memory-associated makers include CD49a (Itga1) and homeobox only protein (hopx), two markers expressed in effector cells and maintained in mCMV--induced memory NK cells (Bezman et al., 2012). During VACV infection, *Itga1* and *hopx* transcripts levels were upregulated in NK cells (Figure 8A). Together, these data suggest that during the acute phase of VACV infection, splenic NK cells substantially upregulate markers that are associated with NK cell memory.

**Figure 8.**
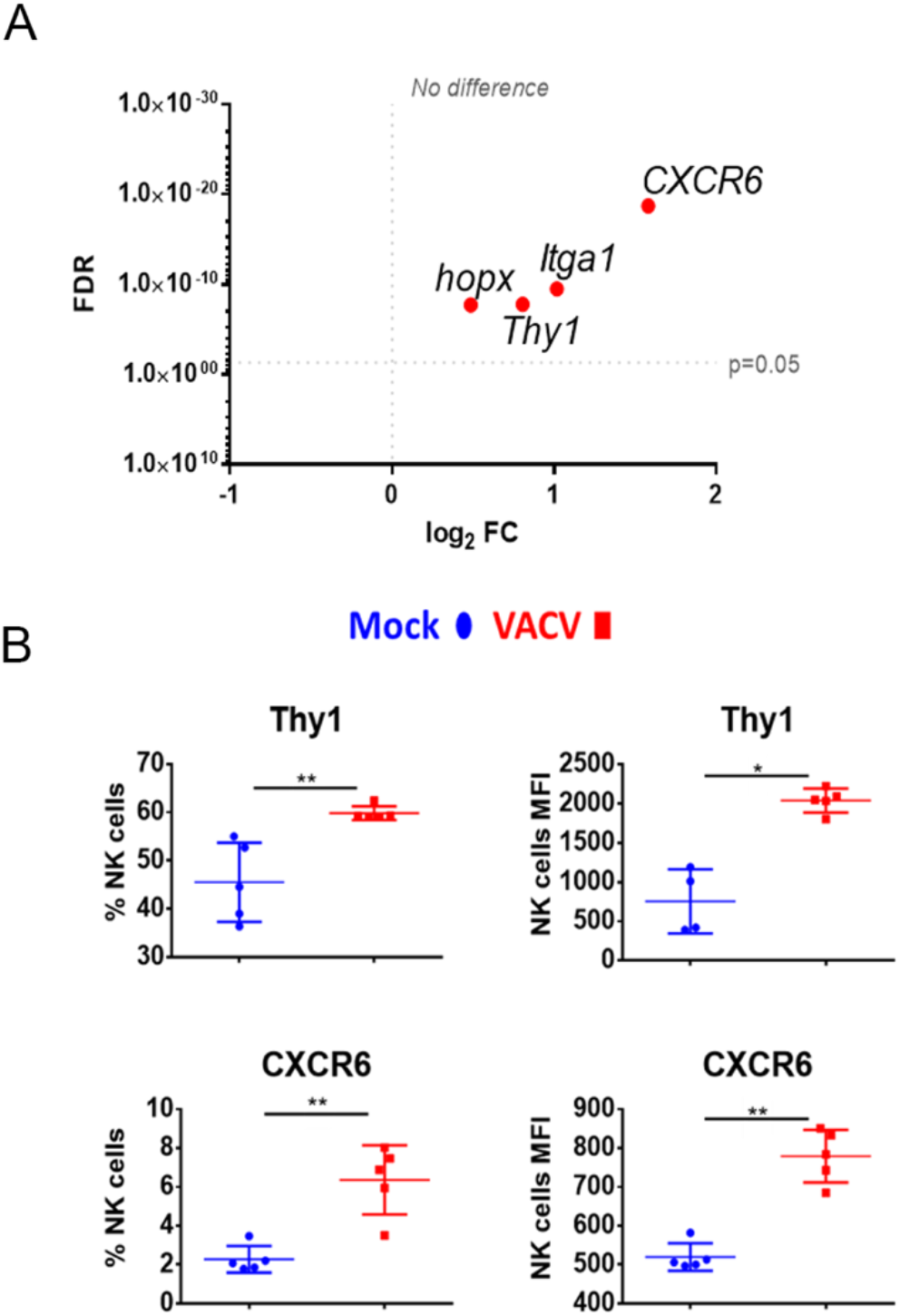
Upregulation of memory NK cell markers, at the transcriptional and protein level during VACV infection. (A) C57BL/6 mice (n=4) were infected with VACV WT or were mock-infected. At 6.5 d.p.i., NK cell transcriptional changes were studied by RNA-seq (n=4), and differential expression (VACV vs mock) was calculated. Each dot represents the differential expression profile of a NK memory marker transcripts (WT vs mock). Red indicates statistical significance (FDR<0.05). For clarity, the protein name is indicated rather than gene name. (B) Protein expression assessed by FACS for SLAM receptors in splenic NK cells from mock (blue) and VACV-infected mice (red). The percentage of NK cells expressing the receptor (prevalence) and MFI (expression level) for the indicated receptor is shown. Error bars represent ± SD, statistical significance was assessed with a Mann-Whitney test (*p<0.05, ** p<0.01, ***p<0.001).

### VACV vaccination induces protective antibodies and CD8+T cells but not protective NK cells

Given the interest in exploiting NK cell memory qualities for vaccination purposes and the observed upregulation of memory markers, we examined whether vaccination with VACV leads to the development of NK cells that can protect from infection. To test this, we infected mice i.d., to mimic vaccination, and four weeks later transferred NK cells, CD8+ T cells or serum from these mice to naïve recipient mice, which were then challenged i.n. with VACV (Figure 9). Both serum and splenic CD8+ T cells conferred some protection against challenge, as shown by reduction in weight loss following infection compared with PBS controls (Figure 9A, B). In contrast, splenic or hepatic NK cells from these immunised mice did not confer protection under these conditions (Figure 9C).

**Figure 9:**
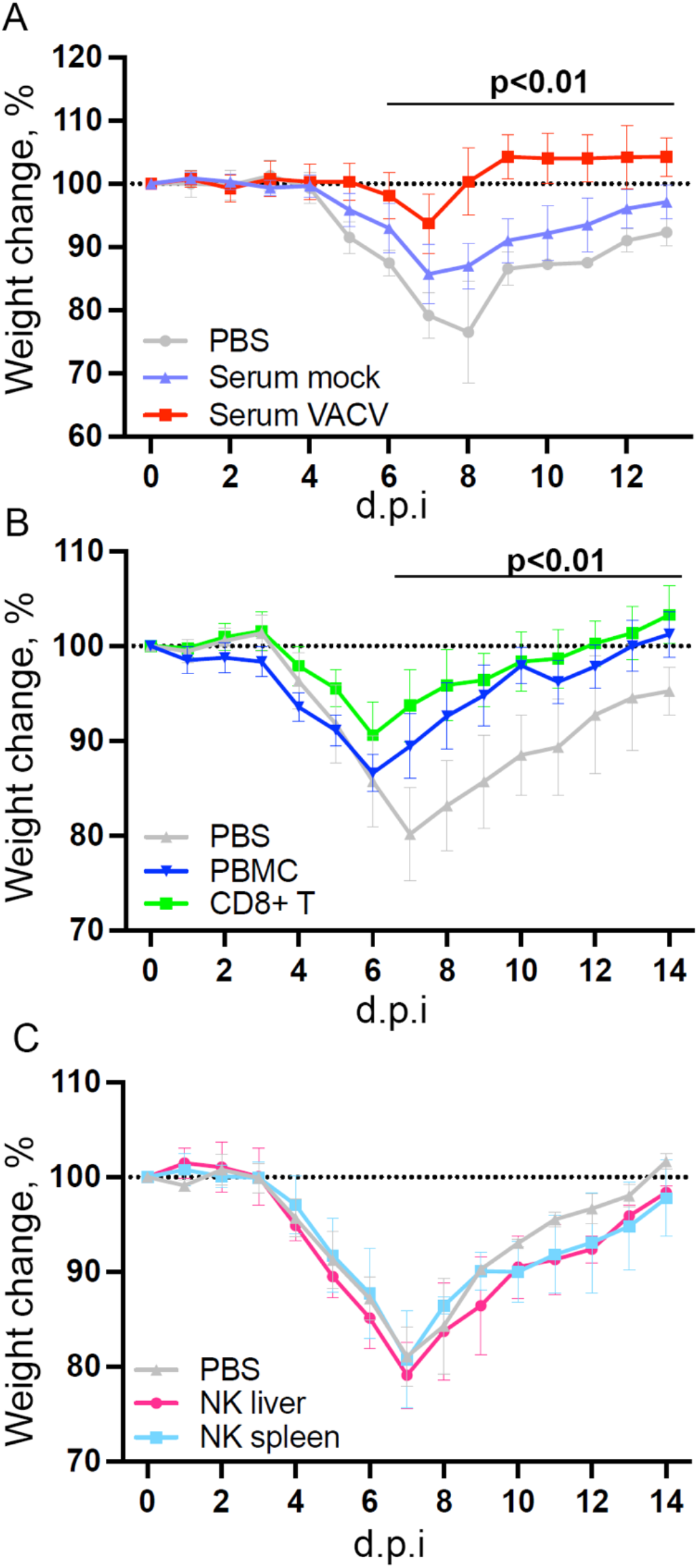
NK cells from vaccinated mice do not confer protection against challenge in a naïve host. C57BL/6 mice were infected i.d. with VACV WR or injected with PBS control and 28 days later (A) Serum was isolated from blood (n=5) or, (B) CD8 T cells (n=10) and PBMC (n=4) or (C) NK cells (n=5) or were purified from spleens. Serum, cells or PBS control (n=4-5) was injected into the tail vein of naïve mice and 24 h later (Day 0) mice were infected i.n. with 10^5^ p.f.u. of VACV WR. Weight change was monitored over the following 14 days.

### VACV and mCMV infection induce NK cell transcriptional programmes with similarities and differences

Next, the NK response to VACV was compared to that of mCMV, another large DNA virus known to generate a robust NK response and to drive memory NK cells development (Smith et al., 2002), with the aim of discovering unique aspects of the NK response to VACV. Using publicly available data (Bezman et al., 2012) we identified NK transcripts that were regulated by one or both viruses over the course of infection in mice (Figure 10). We found that whereas some DEG were changed in response to both viruses, the majority of DEG differed between the two viruses, indicating a unique response to each virus. Genes that were the most upregulated during VACV infection included ISGs, granzymes, serpins, chemokines and their receptors and NK receptors, indicting a more cytolytic activated phenotype of NKs during VACV infection (Figure 10A, B). In contrast, NK gene transcripts uniquely upregulated during mCMV included mainly those associated with regulation of cell cycle and cell division, suggestive of a more proliferative response to mCMV compared to VACV (Figure 10C).

**Figure 10:**
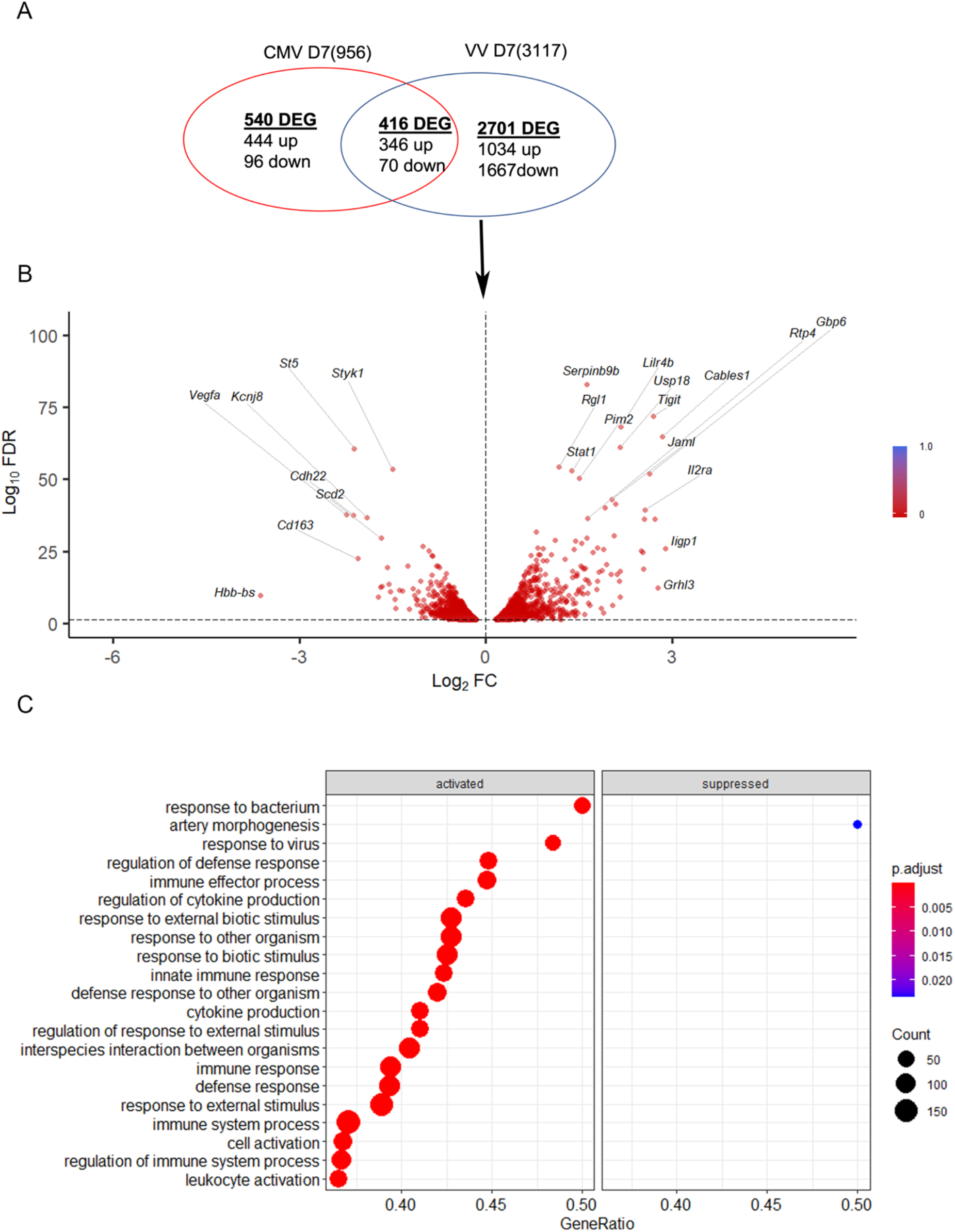
mCMV and VACV induce different gene NK transcriptional profiles in infected mice. (A) Venn diagram showing the number of DEGs (FDR<0.05) (upregulated or downregulated) associated with VACV and mCMV infection in NK cells at d 7 p.i. The total number of genes detected in each dataset is indicated in brackets. (B) Volcano plot of DEG modulated after VACV, but not mCMV infection, in NK cells. (C) Biological processes gene ontologies enriched among the NK cell DEGs modified by VACV infection and not mCMV.

### VACV infection or vaccination induces similar transcription programmes in human and murine NK cells

Finally our transcriptomic data set was compared with a study analysing human NK cells before and 7 d post-vaccination with MVA, an attenuated VACV strain that is used as a vaccine vector and as a vaccine for smallpox and monkeypox (Costanzo et al., 2018). The transcripts of 96 pre-selected genes were studied by qPCR and their FC analysed. Forty-seven transcripts were identified by Costanzo et al. 2018 (FC >1.5 or <-1.5) and thirty-six of these transcripts were detected in our dataset, all of which were described as upregulated in Costanzo et al. (Figure 11). Further, Costanzo and colleagues reported that the transcriptomic changes occurring in NK cells post-MVA-vaccination match closely the transcriptomic signature of NK cells activated by direct recognition. These data are consistent with our observations (Figures 4, 5) and support the view that NK cells from humans and mice share a transcriptomic signature in response to VACV infection or vaccination, which is significantly related to direct recognition of virus-infected cells.

**Figure 11.**
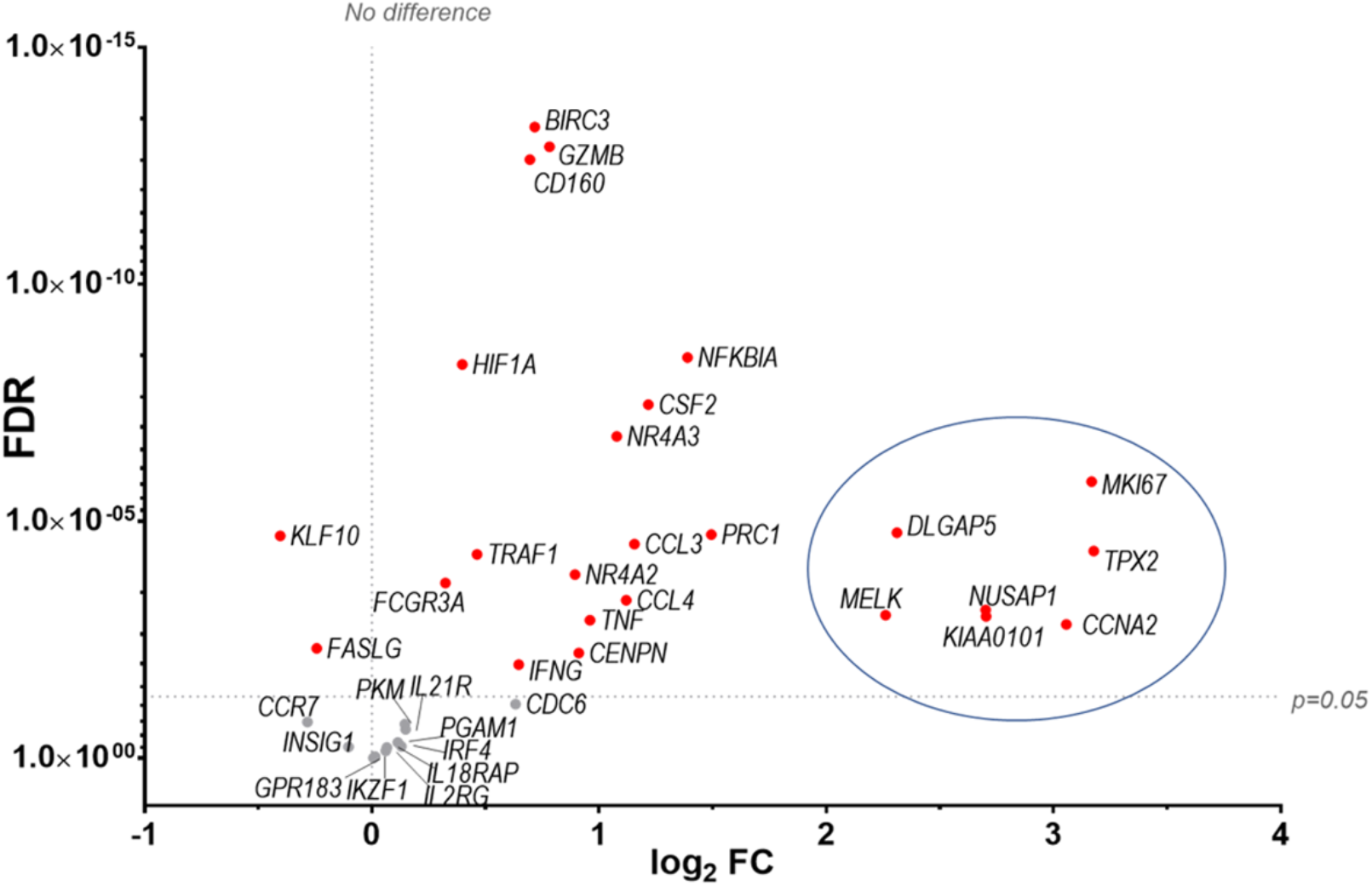
Identification of a transcriptomic signature common to human NK cells post vaccination with MVA and murine NK cells post infection with VACV strain WR. Groups of B6 mice (*n*=4) were mock-infected or infected with VACV WR. At 6.5 d.p.i., NK cell mRNA abundance was studied by RNA-seq and differential expression of transcripts was calculated (WT vs mock). The transcriptomic data (qPCR) from a study looking at human NK cells (n=11) before and at 7 d post MVA vaccination (Costanzo et al., 2018 data reference GSE110446) was compared to our RNA-seq data. Transcripts whose expression post vaccination with MVA was differentially expressed with a FC >±1.5 were selected. The expression of such transcripts was analysed in our RNA-seq data. All such transcripts in our dataset were described as upregulated in Costanzo et al., 2018. Their differential expression in our dataset is represented on the graph. Each dot represents a transcript, the FC (VACV vs mock) is expressed on the X-axis as log2, and the FC statistic value (FDR) is expressed on the Y-axis. Red indicates statistical significance (FDR<0.05). Circled are the transcripts that were most significantly (FDR<0.05) upregulated at 7 d.p.i. with MVA.

## Discussion

NK cells contribute to the clearance of VACV from infected tissues, but there are limited data about how NK cells recognise VACV-infected cells, including which stimuli trigger their activation, which NKR(s) are involved in the recognition and whether they can develop into functional memory cells.

Transcriptomic analysis showed that NK cells undergo broad transcriptional changes in response to VACV infection. DEGs included genes related to immune defence and response to external stimuli, suggesting that NK cells play a role in the immune response to VACV infection. Upon infection, NK cell numbers increased (Figure 1), transcripts for proliferation markers were substantially upregulated and the NK subsets that expanded most are those with cytotoxic and cytolytic capabilities (Figures, 1, 4 and 5) in line with previous reports (Natuk and Welsh, 1986; Dokun et al., 2001; Prlic et al., 2005; Jacobs et al., 2006; Abboud et al., 2016). Additionally, transcripts for mediators of cytotoxic functions (GzmB, GzmK, IFNγ, TNF-α and TRAIL) and protection against damage from cytolytic granules (serpins and cathepsins) were upregulated, indicating that NK cells are equipped to mediate a cytotoxic and cytolytic response. Further, activation markers (CD69, Sca1, KLRG1, CD107a and gp49A/B) were significantly upregulated at the protein and the transcript level suggesting activation. NK memory markers were also upregulated during VACV infection (Figure 8) and given the possibility of VACV-inducing NK memory (Gillard et al., 2011), we explored the ability of NK cells to confer protection against VACV challenge. Using adoptive transfer we showed that serum or memory CD8+ T cells from mice vaccinated one month previously could confer some protection against weight loss induced by i.n. challenge with VACV (Figure 9A, B). However, NK cells purified from the spleen or liver failed to confer protection when transferred into naïve mice that were challenged in the same way (Figure 9C). Previous reports of the protective potential of NK memory-like cells have focussed on their ability to function in immunocompromised mice (RAG1 KO mice) that lack B or T cells. Here we transferred cells into immunocompetent hosts and on this background could not find evidence for their ability to protect from VACV infection.

Comparative analysis with defined NK cell transcriptomic signatures to various stimuli (Costanzo et al., 2018) suggested that NK cells from VACV-infected mice are stimulated primarily by direct target cell recognition. This supports the hypothesis that a VACV-specific NK receptor: ligand couple, similar to mCMV157:Ly49H, could exist and might induce the clonal expansion of a specific NK cell subset. This prompted us to investigate NKRs expression to look for over-represented NKRs that might indicate clonal expansion.

Our results showed stable expression of Ly49H, -D and -G2 at the transcript and protein level, suggesting that these receptors are not likely to define an NK cell subset expanding preferentially during VACV infection. The stable expression of Ly49H is consistent with the literature (Dokun et al., 2001; Gillard et al., 2011), indicating that Ly49H^+^ NK cell expansion during mCMV is specific to this virus. Alternatively, it might reflect the preferential expansion of NK cells that do not express these receptors, similar to the preferential expansion of Ly49H^+^ NK cells, lacking Ly49C/I during mCMV infection in B6 mice (MT et al., 2010). The inhibitory receptors Ly49A and Ly49F were significantly upregulated at the mRNA and protein level but were present as a low percentage of NK cells, 10-15 % and 5-7 % respectively. This suggests that they are unlikely to define an NK cell subset that dominates VACV infection response in a similar way to mCMV-induced expansion of Ly49H^+^ NK cells from 55 % to 90 % at 6.5 d.p.i. (Fogel, 2016). Additionally, Ly49A and Ly49F transcripts were also upregulated in NK cells during mCMV infection, suggesting that their upregulation is not specific to VACV infection.

The study of the SLAM receptor family during VACV infection showed that Ly108 (or NK-T-B-antigen (NTB-A)) and CD319 (or CRACC) were substantially upregulated at the protein and transcript level, whilst other SLAM receptors were not substantially altered. Additionally, the adaptors required for SLAM activating functions (ERT and SAP) were upregulated, suggesting that SLAM receptors were more likely to mediate activating functions. These results deserve further investigation because other SLAM receptors are involved in the NK cell response to various viral infection. For example, human 2B4 and Ly108 bind influenza HA via N-linked glycosylation and lead to NK cell co-stimulation (A et al., 2016) and blocking 2B4 or Ly108 inhibit NK cell-mediated lysis of influenza virus-infected cells (A et al., 2016). Further, soluble decoys for SLAM receptors are found in multiple poxviruses including molluscum contagiosum virus (MOCV) and squirrel poxvirus, which encode orthologues for SLAMF1 and SLAMF2, respectively (Farré et al., 2017).

Comparative analysis showed that five receptors were upregulated substantially during systemic infection with VACV but not with mCMV: NKRP1F, NKRP1B, CRTAM, CD160 and Sema4D. The NKRP1 receptor family and their ligands (CLEC2 subfamily) are of great interest because they are conserved in human and mouse and can mediate surveillance of missing-self (reviewed in (Bartel et al., 2013)). Interestingly, the murine NKRP1B/D ligand, Clr-b, which is expressed ubiquitously, is downregulated by VACV and ectromelia virus (ECTV), which renders target cells more susceptible to lysis by NKRP1B+ NK cells (Williams et al., 2012). Additionally, LLT1 (*clec2d*), the human ligand for human NKRP1A (*klrb1*, inhibitory), which usually is expressed following infection, is downregulated at the mRNA and protein level during VACV infection (Williams et al., 2016). Moreover, another unidentified protein is upregulated during VACV WR infection, cross-links with the 4C7 mAb (raised against clec2d), and has an expression kinetic that is inversely correlated with the degradation of clec2d mRNA (Williams et al., 2016). These data suggest that during VACV infection, a decoy ligand might be expressed to replace loss of clec2d. Further, CD160 in human and mice binds classical and non-classical MHC-I (Barakonyi et al., 2004; Maeda et al., 2005), but also HVEM (herpesvirus entry mediator) and enhances NK cell lytic activity and cytokine secretion (Le Bouteiller et al., 2011; Šedý et al., 2013; Tu et al., 2015; Liu et al., 2019). Interestingly, CD160 upregulation was also observed in human NK cells post MVA vaccination (Costanzo et al., 2018), suggesting that CD160 might be involved in the recognition of poxvirus-infected cells, and that it could be conserved in mouse and humans. Hence, CD160 is a strong candidate receptor to investigate further in the context of VACV infection.

In summary, this FACS study of NKR expression during VACV infection did not highlight a dominating NKR similar to that observed during mCMV infection. However, in combination with transcriptomic and bioinformatic studies, this allowed the identification of a few candidate NKRs (Ly108, CD319, NKRP1B, NKRP1F, CD160) that may be involved in the immune response to VACV. Markers known to be associated with the development of memory NK cells were upregulated after VACV infection at 6.5 d.p.i., suggesting that a subset of circulating NK could retain memory qualities and remain long-lived. However, the adoptive transfer of splenic or hepatic NK cells from vaccinated mice into naïve mice did not confer protection upon VACV challenge. On the contrary, the adoptive transfer of serum, CD8+ T cells and total splenocytes did confer protection upon challenge. This is consistent with the established ability of VACV to induce the development of strong and specific humoral and CD8+ memory cells. The inability of VACV-primed NK cells to confer protection is in contrast with a study reporting that Thy1+ hepatic VACV-primed NK cells that were adoptively transferred into naïve into RAG1ko mice conferred protection upon viral challenge (Gillard et al., 2011). Multiple reasons could explain this discrepancy. The main difference is the recipient mice used. Whilst we used immunocompetent mice, Gillard et al used Rag1 knockout mice, in which transferred NK cells may provide their protective capacities in a niche now unoccupied by other lymphocytes. In an extension to this argument, multiple studies which demonstrated NK memory protective abilities were performed in new-born or immunodeficient mice (Paust et al., 2010; Gillard et al., 2011). Alternatively, immunodeficient recipient might allow for higher amplification and better survival of transferred NK cells than WT hosts.

Collectively, our study provides new data about NK cell function and homeostasis during VACV infection that has implications for the design of poxvirus-based vaccines for heterologous pathogens and VACV-based oncolytic therapy and further our understanding of NK cell biology and host-pathogen interactions.

## Author Contributions

BJF, DD, and GLS contributed to conception and design of the study. DD performed the experiments. BJF, DD, and GLS analysed the data. BJF, DD, and GLS drafted, read, and approved the submitted manuscript

## Acknowledgements

This work was supported by UKRI MRC grant MR/M019810/1 to GLS and BF. The authors declare no conflicts of interest. We thank the Phenotyping hub flow cytometry facility, University of Cambridge, UK for assistance with flow cytometry cell sorting, Cambridge Genomics Services for assistance with RNAseq analysis and library preparation, and the Pathology animal facility staff for support with animal husbandry and studies. We also thank Dr Zhenya Shmeleva for performing intravenous injections in adoptive transfer experiments and Dr Franceso Colucci for helpful discussion on the dataset.

## Data Availability Statement

Datasets for this study can be found in the Europen Nucleotide Archive https://www.ebi.ac.uk/ena, PRJEB57353

## Ethics statement

Animal experiments in the UK were conducted according to the Animals (Scientific Procedures) Act 1986 under PPL 70/8524 issued by the UK Home Office.

## Supplementary Data

**Supplementary Figure 1.**
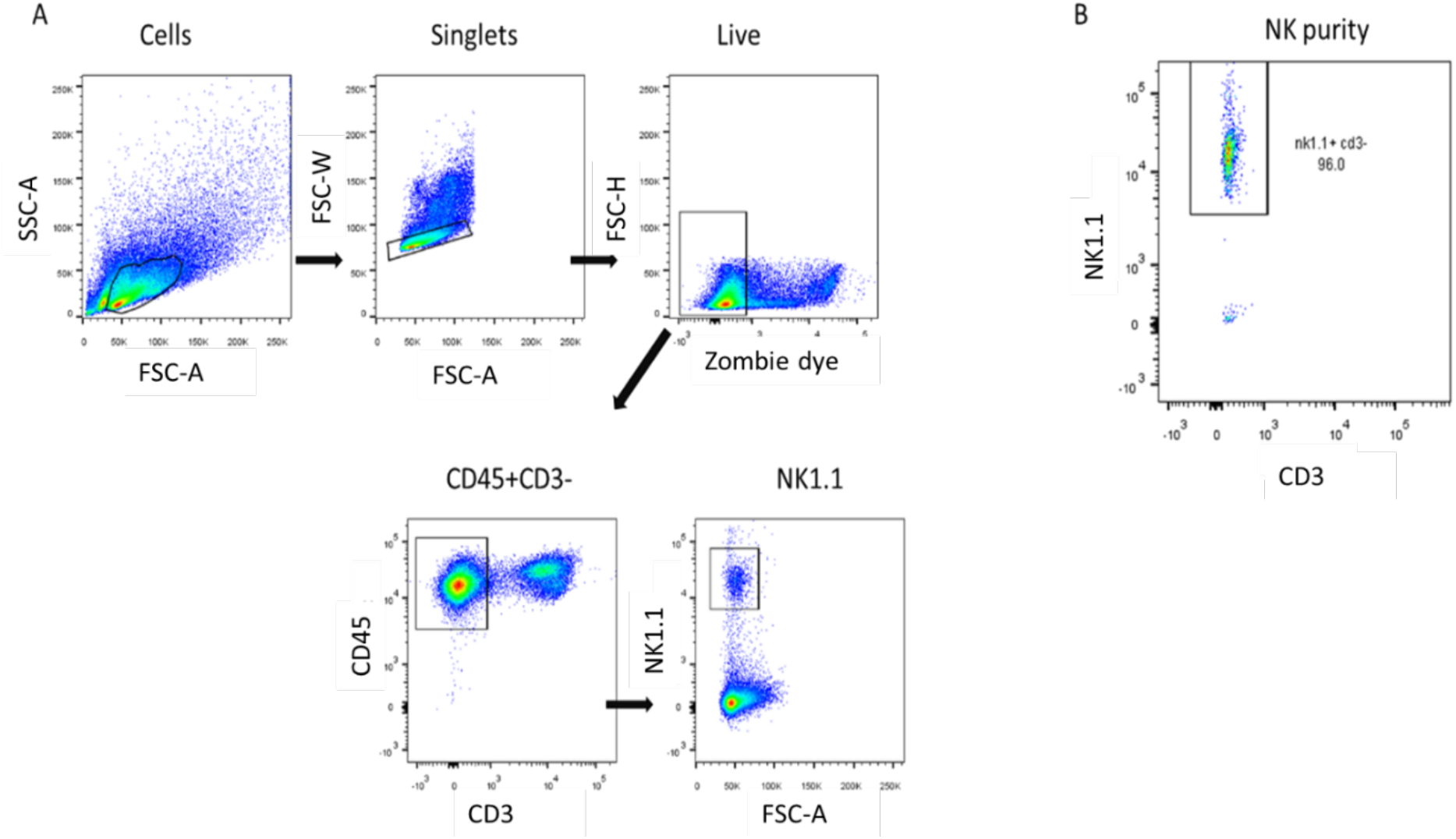
Gating strategy for NK cell isolation by FACS and purity check. (A) Debris were gated out and viable single cells that were negative for CD3, and positive for CD45 and NK1.1, were isolated by FACS in sterile PBS. (B) The purity of the NK cells was checked by running a sample of the sorted cells on the flow cytometer.

**Supplementary Figure 2.**
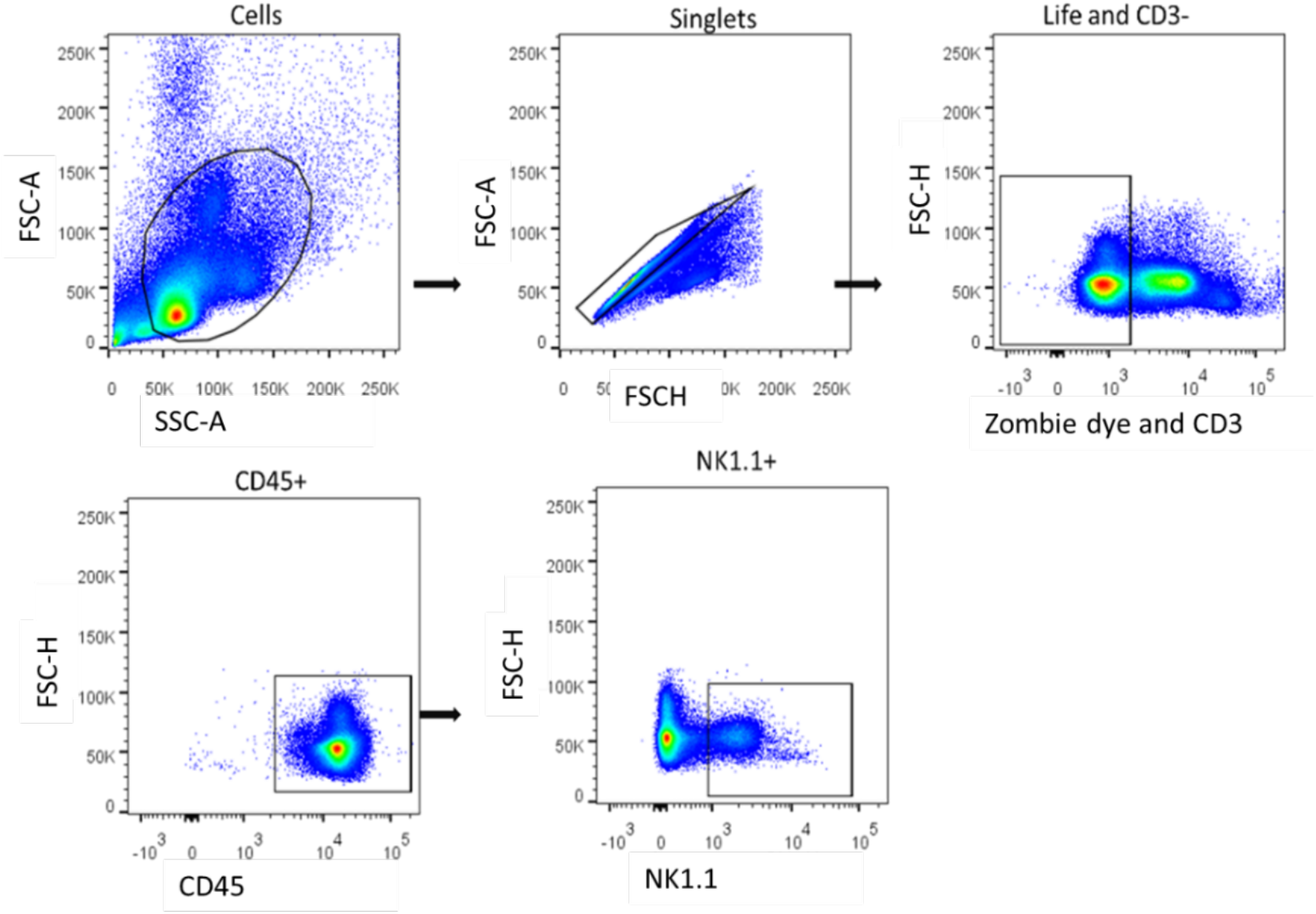
Common gating strategy used for NK phenotyping by FACS. Debris were gated out and then viable single cells negative for CD3, and positive for CD45 and NK1.1 were gated.

